# Transcriptomic Integration Reveals a Conserved Inflammatory–Proliferative Paradox in Acquired Resistance to Immune Checkpoint Blockade

**DOI:** 10.64898/2026.04.01.714095

**Authors:** Hoyeon Lee, Heerim Yeo, Inseon Bak, Kyeong-Won Yoo, Sang-Min Park

## Abstract

Acquired resistance to immune checkpoint blockade (ICB) is increasingly recognized as an active adaptive process. However, prior studies have typically focused on individual tumor models, limiting the ability to distinguish conserved mechanisms from model-specific observations. Here, we integrated four independent transcriptomic datasets of acquired ICB resistance, spanning human non-small cell lung cancer (NSCLC) biopsies, murine CT26 colorectal tumors, organoid-derived murine NSCLC tumors, and EMT6 breast cancer cells. Differential expression analysis was performed within each dataset, followed by an intersection-based consensus approach to identify reproducible resistance-associated programs. Contrary to the conventional cold tumor paradigm, acquired-resistance tumors consistently maintained interferon-γ response and innate immune signaling while simultaneously activating cell-cycle programs and constitutive KRAS signaling signatures across all four models. We term this an apparent inflammatory–proliferative paradox: the persistence of IFN-γ–driven inflammatory signatures, canonically associated with productive antitumor immunity, in tumors that have escaped immune control. Notably, this program was retained in immune-depleted organoid and cell-line models, supporting a tumor-cell-associated component maintained independently of the immediate immune microenvironment. Transcription factor activity inference identified a conserved regulatory backbone linking interferon-associated regulators (*STAT2*, *IRF2*) with proliferation drivers (*E2F4*, *TFDP1*) and suppression of lineage-specifying factors (*HNF4A*, *EGR1*). Integrated network analysis resolved these signals into three reinforcing modules, namely hyper-proliferative outgrowth, active inflammatory adaptation, and lineage identity loss. This architecture provides a systems-level framework for prioritizing combination strategies that simultaneously address interconnected resistance axes.

## Introduction

Over the past decade, the development of immune checkpoint inhibitors (ICIs) has fundamentally transformed the therapeutic landscape of solid tumors [1]. In particular, antibody therapies targeting programmed death-1 (PD-1) and programmed death-ligand 1 (PD-L1) have established a new treatment paradigm by restoring impaired antitumor immunity rather than directly targeting tumor cells, thereby distinguishing themselves from conventional chemotherapy and targeted therapies [2]. PD-1 blockade has demonstrated significant survival benefits across multiple solid malignancies, including melanoma, lung cancer, renal cell carcinoma, and head and neck cancers [3].

Despite these advances, the overall response rate to ICIs remains limited at approximately 20–30% [4], and even patients who initially respond frequently experience disease recurrence or progression due to acquired resistance. Moreover, a subset of patients derives no clinical benefit from the outset of treatment, a phenomenon known as primary resistance [5]. These resistance patterns highlight the complex and multifactorial biological processes that underlie immunotherapy failure [6]. Thus, defining the molecular mechanisms that give rise to acquired resistance is critical for overcoming current barriers in cancer immunotherapy.

More broadly, cancer is a highly heterogeneous disease shaped by inter-patient variability, disease stage, and the anatomical context in which tumors arise [7]. Even within the same patient, tumors located in different tissue or metastatic sites can exhibit distinct cellular compositions and immune microenvironments [8]. Consequently, cancers sharing the same histological diagnosis may behave as biologically distinct entities in their interactions with the immune system [9]. This context-dependent heterogeneity is increasingly recognized as a major determinant of therapeutic outcomes and provides a conceptual basis for the heterogeneous responses observed following immune checkpoint blockade.

One widely used framework to explain heterogeneous ICI responses is the concept of immunologically “hot” versus “cold” tumors, defined by the degree of T-cell infiltration and interferon-γ (IFN-γ)–driven inflammation within the tumor microenvironment (TME) [10]. Hot, inflamed tumors with abundant CD8_ T cells and IFN-γ signaling are generally more responsive to PD-1/PD-L1 blockade, whereas cold tumors with sparse immune infiltrates and immunosuppressive myeloid populations often show primary resistance. However, this binary classification does not fully capture clinical reality. Patients with non-small cell lung cancer (NSCLC) who have developed acquired resistance to immunotherapy can display paradoxically increased IFN-γ signatures, while these signals no longer translate into effective tumor clearance [11]. Similarly, in an EMT6 breast cancer model, chronic PD-L1 inhibition leads to a “hot but dysfunctional” phenotype characterized by sustained type I IFN signaling with a secretory program that protects tumor cells from immune attack [12]. In addition, acquired resistance can also arise through tumor cell–intrinsic upregulation of collagen and TGF-β–driven extracellular matrix remodeling, which creates physical barriers that exclude T cells from the tumor core [13]. Collectively, these studies support a view of acquired resistance as an active adaptive process; however, most rely on a single tumor type or experimental system. Relying on a single experimental system poses a risk of capturing model-specific artifacts rather than universal mechanisms. For instance, syngeneic mouse models may not fully reflect human immune complexity, while clinical biopsies are often confounded by high inter-patient heterogeneity. While chronic inflammatory signaling has been individually implicated in immune dysfunction and therapy resistance, whether it co-occurs with proliferative activation and lineage identity loss as part of a conserved cross-model program remains unexplored.

To bridge this gap, we developed a cross-species, cross-platform transcriptomic framework using four independent models of acquired resistance to anti–PD-1/PD-L1 therapy: human NSCLC biopsies, murine CT26 colorectal cancer, murine NSCLC organoids, and EMT6 murine breast cancer cells. Together, these datasets span *in vivo* tissues and immune-depleted *in vitro* conditions, enabling us to capture conserved tumor-cell-associated adaptations beyond any single tumor type or platform. Using pathway enrichment analysis, transcription factor inference, and integration of recurrently altered genes, we investigated whether acquired resistance converges on a unified program linking persistent inflammatory signaling, proliferative activation, and lineage identity loss. By defining this interconnected resistance program and its regulatory backbone, we seek to provide a systems-level framework for understanding ICI failure and to nominate candidate molecular targets for preventing or overcoming acquired resistance across diverse tumor contexts.

## Materials and Methods

### Data acquisition

Transcriptomic datasets modeling acquired resistance to immune checkpoint blockade were obtained from the Gene Expression Omnibus (GEO) database. We included one human microarray dataset and three murine-derived RNA-seq datasets spanning a syngeneic tumor model, an organoid model, and a cell-line model: H-NSCLC (GSE248249), M-CRC (GSE249000), M-NSCLC (GSE261889), and M-BRCA (GSE186034).

The human dataset, H-NSCLC (GSE248249), comprises microarray profiles of tumor biopsy samples from patients with NSCLC treated with anti–PD-1 therapy [11]. The dataset includes pretreatment samples collected prior to therapy (Pre, n = 13) and post-treatment samples collected at the time of clinically confirmed disease progression following initial response (Post, n = 29). Biopsies originated from both primary and metastatic sites.

For murine models, M-CRC (GSE249000) is an RNA-seq dataset generated from a syngeneic CT26 murine colorectal cancer model [11]. Samples include parental CT26 tumors prior to anti–PD-1 treatment (CPar, n = 3) and tumors that acquired resistance following iterative *in vivo* anti–PD-1 selection. Resistant tumors were generated through repeated cycles of tumor implantation, anti–PD-1 treatment, tumor excision, and reimplantation, yielding second-generation (C2nd, n = 2) and fourth-generation (C4th, n = 4) resistant tumors. Tumor-derived cells were expanded *ex vivo* prior to RNA-seq analysis.

M-NSCLC (GSE261889) is an RNA-seq dataset derived from a genetically engineered murine NSCLC organoid model with *Trp53* loss, *Kras*^G12D activation, and *Myc* overexpression [13]. Following orthotopic transplantation into immunocompetent mice, tumors were treated with anti–PD-1 therapy and classified as pretreatment tumors (NSCLC, n = 4), anti–PD-1–sensitive tumors that remained responsive to treatment (Sen, n = 3), or tumors that acquired resistance after prolonged therapy (Res, n = 9). RNA-seq was performed after *in vivo* selection followed by *ex vivo* organoid culture to enrich for tumor-cell-associated transcriptional programs.

Finally, M-BRCA (GSE186034) is an *in vitro* RNA-seq dataset derived from the EMT6 murine breast cancer cell line [12]. Cells were treated continuously with anti–PD-L1 antibody for more than four weeks, generating PD-L1–tolerant cells (PTR, n = 3) and matched parental controls (P, n = 3). This dataset represents tumor-intrinsic transcriptional adaptations to chronic PD-L1 blockade in the absence of immune and stromal components.

### Gene expression analysis

Bioinformatics analysis was performed in R (v4.4.3). Raw count data for RNA-seq datasets (M-CRC, M-NSCLC, and M-BRCA) were analyzed using *DESeq2* (v1.46.0), whereas normalized microarray intensities for H-NSCLC were analyzed using *limma* (v3.62.2). Principal component analysis (PCA) was conducted independently for each dataset to visualize global transcriptomic shifts associated with resistance. To avoid technical artifacts from cross-platform normalization (microarray versus RNA-seq), all differential expression and downstream enrichment analyses were conducted independently within each dataset to generate dataset-specific statistics. For each dataset, comparisons contrasted the resistant state against the sensitive or parental baseline: Post vs Pre (H-NSCLC), C4th vs CPar (M-CRC), Res vs NSCLC (M-NSCLC), and PTR vs P (M-BRCA). For RNA-seq analysis, size-factor normalization and dispersion estimation were performed using the default DESeq2 workflow. Differential expression was assessed using the Wald test, with p-values adjusted for multiple testing using the Benjamini–Hochberg false discovery rate (FDR) procedure. For the H-NSCLC microarray dataset, differential expression between Pre- and Post-treatment samples was modeled using limma. Conserved molecular features were subsequently identified using an intersection-based cross-dataset consensus approach, selecting signatures that were statistically significant within individual datasets and exhibited concordant directionality across all models. Mouse-to-human gene ortholog information was retrieved from the National Center for Biotechnology Information (NCBI).

### Pathway enrichment analysis

Pathway analysis was performed using gene set enrichment analysis (GSEA) and over-representation analysis (ORA). GSEA was performed via the fgsea package (v1.32.4), with genes ranked by differential expression statistics: Wald statistics from DESeq2 for murine datasets and moderated t-statistics from limma for the human dataset. For microarray data with multiple probes per gene, the probe with the maximum absolute statistic was selected. Gene sets were retrieved from the Molecular Signatures Database (MSigDB) via msigdbr (v25.1.1), encompassing Hallmark (H/MH), Gene Ontology (GO), BioCarta, Reactome, and WikiPathways collections. Species-specific collections (C2/C5 for human and M2/M5 for mouse) were utilized, filtering for gene sets containing 5–500 genes. Statistical significance was estimated using 100,000 permutations, with pathways considered significant at a false discovery rate (FDR) < 0.05. For ORA, upregulated and downregulated genes were analyzed respectively using the Enrichr web server to identify over-represented biological pathways [14]. Statistical significance was assessed via the hypergeometric test, with Benjamini–Hochberg adjusted P values < 0.05 considered significant. To assess the overlap in gene composition among the three pathways consistently regulated across all four datasets, Pairwise Jaccard indices were calculated as the ratio of the intersection size to the union size of the gene sets.

### Transcription factor analysis

To identify candidate regulatory drivers underlying these transcriptomic changes, transcription factor (TF) activity scores were inferred using *decoupleR* (v2.8.0) [15]. For each dataset, differential expression statistics from the resistant-versus-baseline contrast were used as the input signature. Specifically, the limma moderated t statistics (H-NSCLC) and DESeq2 Wald statistics (M-CRC, M-NSCLC, and M-BRCA) were supplied to *decoupleR*. TF activity scores were computed using the *run_viper* function, which implements the VIPER (Virtual Inference of Protein Activity by Enriched Regulon Analysis) algorithm, to estimate protein-level activity based on the expression of downstream target genes [16]. Species-matched DoRothEA regulons were used for human (H-NSCLC) and mouse datasets (M-CRC, M-NSCLC, and M-BRCA).

### Network construction and visualization

Integrated networks were constructed in Cytoscape (v3.10.3) by combining the identified TFs, DEGs, and enriched pathways from GSEA and ORA. Edges were defined based on two criteria. TF–DEG interactions were established using species-matched DoRothEA regulons, while pathway–gene associations were determined by gene-set membership. Specifically, pathway nodes derived from GSEA were linked to all matching TFs and DEGs within the corresponding MSigDB gene set, whereas nodes derived from ORA were connected exclusively to the subset of DEGs mapped to that term. For visualization, distinct shapes were assigned to differentiate TFs, DEGs, and pathways. TF and DEG nodes were sized proportional to their degree and colored according to activity scores and fold-change values, respectively, while pathway nodes were assigned discrete colors based on functional module. Module assignment was performed by manual annotation based on the functional coherence of constituent nodes. Pathways, TFs, and DEGs were grouped according to their established biological roles in proliferation, inflammatory/immune signaling, or lineage specification.

### Network module calculation

To evaluate the structural properties of each module, network topology analysis was performed by quantifying internal and external connectivity. Internal edges (£_in_) were defined as connections between nodes within the same module, while external edges (£_out_) represented connections between nodes in the module and nodes outside the module. For a module containing n nodes in a network with a total of N nodes, the number of possible internal edges was calculated as n(n−1)/2. The number of possible external edges was calculated as n(N−n), representing the total potential connections between the n nodes within the module and the (N−n) nodes outside it. Internal density was defined as the proportion of observed internal edges among all possible internal edges, £_in_/[n(n−1)/2], whereas external density was defined as the proportion of observed external edges among all possible external edges, £_out_/[n(N−n)]. These metrics were used to assess the degree of topological cohesion within each module and its relative connectivity to the rest of the network.

## Results

### Dataset overview and global transcriptomic structure

To identify conserved mechanisms of acquired resistance to immune checkpoint blockade, we performed an integrated transcriptomic analysis of four independent datasets derived from diverse biological models (**Fig. 1A**). These included (i) human non-small cell lung cancer (NSCLC) biopsies obtained at baseline (pre) and at disease progression (post) following anti-PD-1 treatment [11], (ii) CT26 murine colorectal tumors that developed resistance following iterative *in vivo* anti-PD-1 treatment and reimplantation [11], (iii) murine NSCLC organoids selected for resistance under anti-PD-1 pressure in immunocompetent hosts and then profiled *ex vivo* to focus on tumor-cell-associated changes, and (iv) EMT6 murine breast cancer cells generated through continuous *in vitro* exposure to anti-PD-L1 antibody, representing a resistance phenotype evolved in the absence of immune cell interaction. Together, these four systems span species (mouse and human), platforms (RNA-seq and microarray), tissue origins (lung, colon, and breast), and experimental contexts (*in vivo* tissues, *ex vivo* organoids, and *in vitro* cell lines), while sharing a common pre-treatment versus acquired resistance structure (**Table 1**).

**Figure 1.**
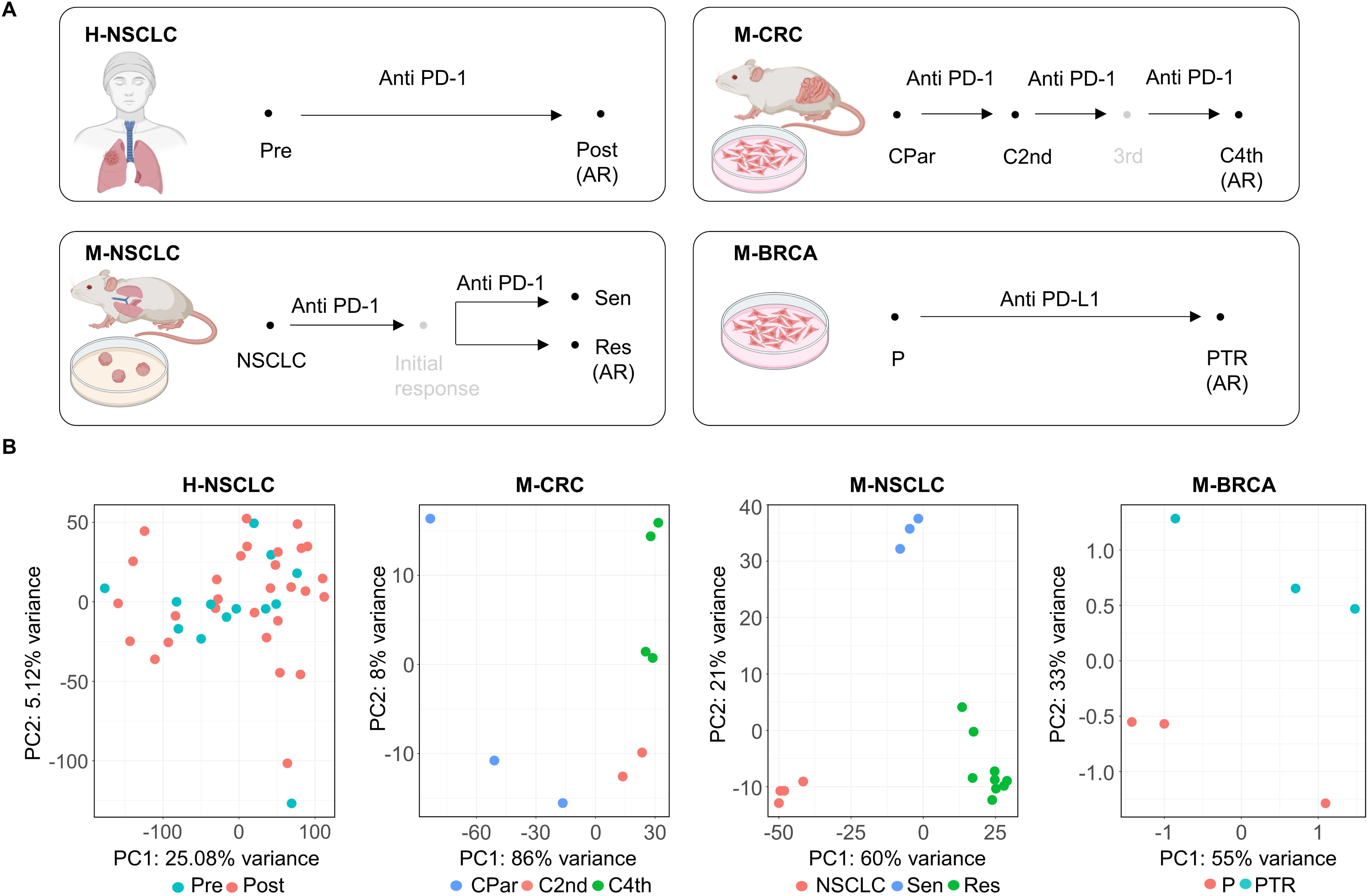
Study design and global transcriptomic landscape of acquired resistance models. **(A)** Schematic illustration of the four independent acquired resistance models analyzed in this study. H- NSCLC represents human non-small cell lung cancer biopsies collected at baseline (Pre) and at the time of disease progression (Post) following anti-PD-1 therapy. M-CRC represents syngeneic CT26 murine colorectal tumors evolved through iterative *in vivo* anti-PD-1 selection, profiled as parental (CPar), second-generation (C2nd), and fourth-generation (C4th) tumors. M-NSCLC represents genetically engineered murine lung cancer organoid-derived model following *in vivo* anti–PD-1 treatment, including baseline tumors (NSCLC), anti–PD-1–sensitive tumors (Sen), and resistant tumors (Res). M-BRCA represents EMT6 murine breast cancer cells generated through chronic *in vitro* exposure to anti-PD-L1 antibody, profiled as parental (P) and resistant (PTR) cells. AR indicates the acquired resistant state in each model. **(B)** Principal component analysis (PCA) performed independently within each dataset after dataset-specific normalization. Percent variance explained by PC1 and PC2 is indicated on each axis.

**Table 1.** Overview of transcriptomic datasets and biological models of acquired resistance to PD-1/PD-L1 blockade used in this study.

| GEO<br>accession | Species | Cancer Type | Sample Type | Drug | Group | N | Platform |
| --- | --- | --- | --- | --- | --- | --- | --- |
| GSE248249 | Human | NSCLC | Tumor biopsy | Anti-human PD-1 | Pre (Treatment-naive),<br>Post (Resistant) | Pre: 13<br>Post: 29 | Microarray |
| GSE249000 | Mouse<br>(BALB/c) | Colorectal cancer (CT26) | Tumor biopsy<br>(subcutaneous) | Anti-mouse PD-1 | CPar (Parental),<br>C2nd (Intermediate),<br>C4th (Resistant) | CPar: 3<br>C2nd: 2<br>C4th: 4 | RNA-seq |
| GSE261889 | Mouse<br>(C57BL/6) | Mouse lung organoids with Trp53<br>mutation, Kras mutation, and Myc<br>amplification (TKM) | Tumor biopsy (orthotopic) | Anti-mouse PD-1 | NSCLC,<br>Res (Resistant),<br>Sen (Sensitive) | NSCLC: 4<br>Res: 9<br>Sen: 3 | RNA-seq |
| GSE186034 | Mouse<br>(BALB/c) | Breast cancer (EMT6) | Cell line (*In vitro*) | Anti-mouse PD-L1 | P (Parental),<br>PTR (Resistant) | P:3<br>PTR:3 | RNA-seq |

After dataset-specific normalization, principal component analysis (PCA) was performed separately for each model. These analyses confirmed that all four datasets capture substantial transcriptional reprogramming associated with acquired resistance (**Fig. 1B**). While isogenic murine models exhibited clear global separation between sensitive and resistant tumors, human clinical samples showed significant overlap with only a more gradual but consistent shift from ‘pre’ to ‘post’, reflecting high inter-patient heterogeneity. This discrepancy highlights the challenge of detecting resistance drivers in clinical datasets and underscores the necessity of a pathway-based meta-analysis to identify conserved molecular mechanisms that transcend individual model- or patient-specific variability. Consistent with the PCA results, volcano plots for each cohort revealed widespread differential gene expression between sensitive and resistant states, although the extent and distribution of transcriptional changes varied across models (**Fig. S1**). These findings further support the presence of broad transcriptomic remodeling across all four resistance models.

### Pathway-level analysis reveals a shared inflammatory–proliferative program

To uncover the functional convergence underlying acquired resistance beyond individual gene variations, we performed Gene Set Enrichment Analysis (GSEA) across four distinct models. Remarkably, among all 13,563 gene sets analyzed, only three pathways exhibited consistent and significant enrichment across all four datasets (**Fig. 2A**). We observed a universal upregulation of inflammatory signaling pathways across all four models. The “Interferon Gamma Response” and “Regulation of Innate Immune Response” gene sets were consistently enriched in resistant samples compared to their sensitive counterparts. Importantly, genes associated with the IFN-γ response pathway and innate immune response pathway remained consistently enriched not only in immune-competent tissue models (H-NSCLC, M-CRC) but also in models devoid of immune cells, including organoids (M-NSCLC) and *in vitro* cells (M-BRCA). This observation suggests that these inflammatory response-associated transcriptional features in acquired resistance are not solely driven by immune cell infiltration but may include a stable tumor-cell-associated component. In addition, we observed consistent enrichment of the “Neoplastic Transformation KRAS Up” signature, a transcriptional marker of constitutive KRAS signaling. This suggests that resistant tumors adopt a sustained, KRAS-like proliferative program to withstand immune pressure. Pairwise overlap analysis showed minimal gene sharing among the three pathways (Jaccard indices = 0.01–0.07; **Fig. S2**), indicating that their identification reflects independent biological convergence rather than redundancy in gene-set composition.

**Figure 2.**
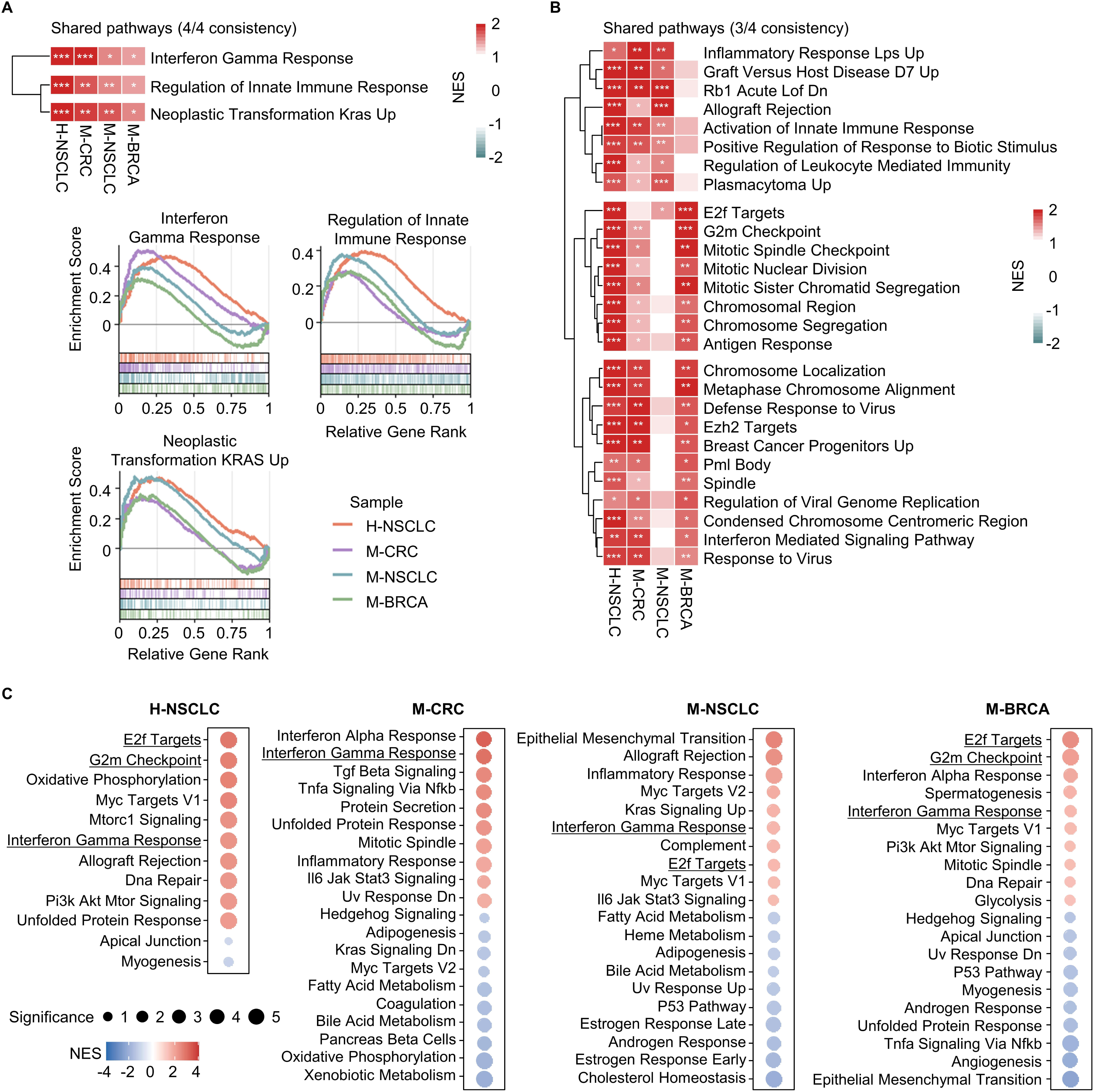
Pathway-level analysis using shared signatures. **(A)** Heatmap of normalized enrichment scores (NES) showing pathways commonly shared across all four datasets (top), and representative GSEA enrichment plots for the same shared pathways (bottom). *FDR < 0.05, **FDR < 0.01, ***FDR < 0.001. **(B)** Heatmap of normalized enrichment scores (NES) showing pathways commonly shared across three datasets. **(C)** Dot plots summarizing the top and bottom Hallmark pathways for each cohort, ranked by NES. Dot color represents NES, and dot size indicates pathway significance (- log_10_ *P-*value).

When we relaxed the criterion to pathways enriched in three of four models, an extended set of shared signatures emerged (**Fig. 2B**). These reinforced the dual nature of resistance, featuring additional inflammatory pathways (“Inflammatory Response,” “Defense Response to Virus,” “Regulation of Leukocyte Mediated Immunity”) together with multiple cell cycle modules (“E2F Targets,” “G2M Checkpoint,” “Mitotic Spindle”, “Chromosome Segregation”). Consistent with the shared pathway analysis, cohort-specific dot plots of Hallmark pathways revealed that inflammatory signaling and cell cycle programs were among the most strongly enriched features in resistant samples across all models (**Fig. 2C**), supporting the robustness of the inflammatory–proliferative program at the individual cohort level.

Thus, across species and experimental contexts, tumors with acquired resistance consistently combined sustained immune signaling with activation of proliferative programs. Collectively, these pathway-level results define a shared “inflammatory–proliferative paradox”: tumors that have escaped PD-1/PD-L1 therapy remain transcriptionally inflamed yet simultaneously reinforce cell cycle and mitotic machinery, supporting continued growth despite ongoing immune pressure.

### Transcription factor analysis characterizes convergent reprogramming driving the inflammatory–proliferative phenotype

We next sought to delineate the regulatory network underlying the shared phenotype of activated immune signaling and concurrent cellular growth in acquired resistance by examining transcription factor (TF) activity patterns across the four independent models. We first assessed the intersection of significantly altered TFs showing consistent directionality across samples to identify a universal regulatory factor (**Fig. 3A**). The initial intersection analysis revealed substantial heterogeneity among models, with no single TF universally regulated across all four datasets.

**Figure 3.**
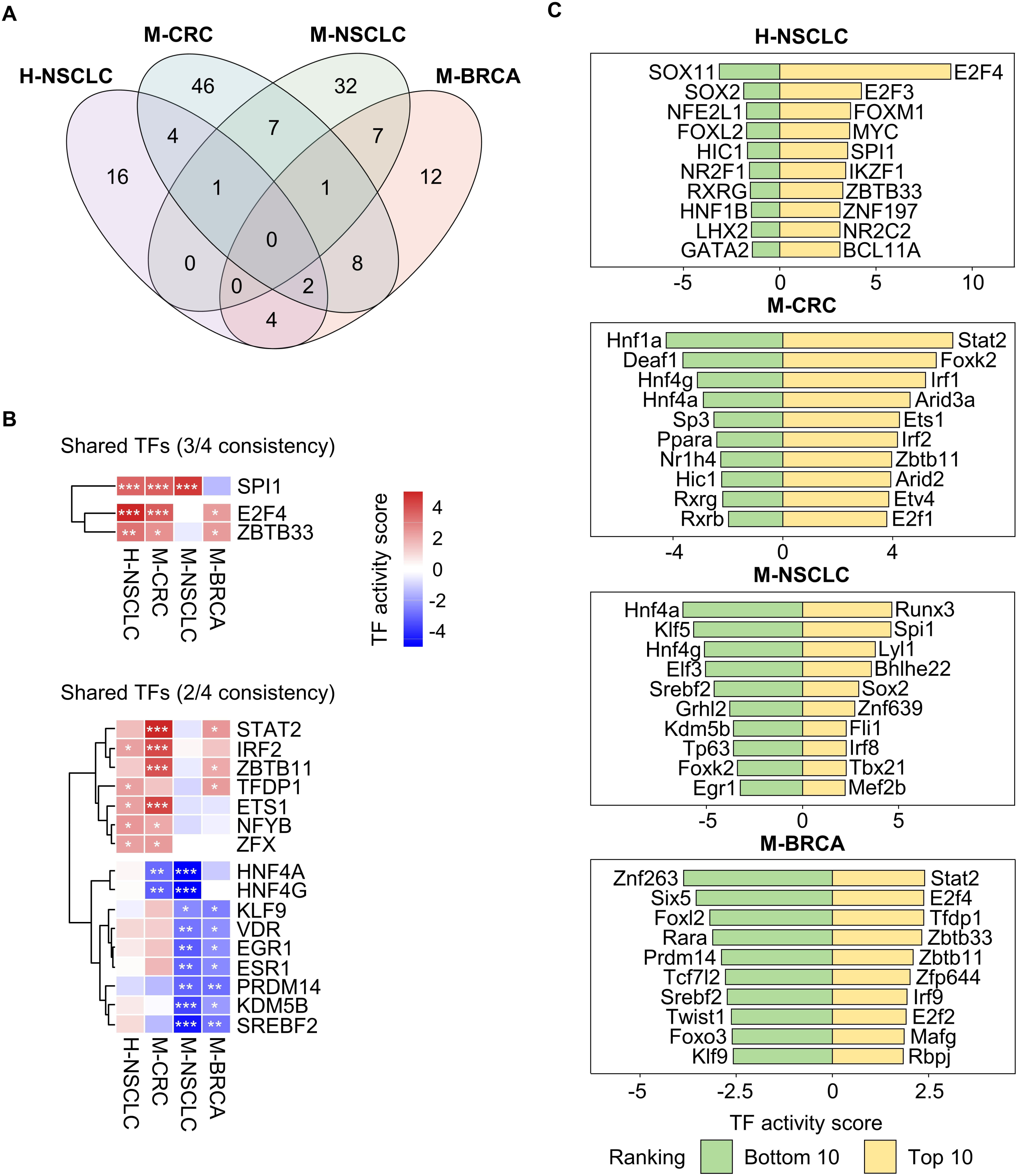
Identification of shared transcriptional drivers across resistant tumors. **(A)** Intersection analysis of significantly altered transcription factors (TF) across the four independent resistant models (H-NSCLC, M-NSCLC, M-CRC, and M-BRCA). **(B)** Conserved TF signatures identified by focusing on factors consistently altered in at least three (top) or two (bottom) models. The heatmap visualization displays TF activity scores. \**P* <0.05, \*\**P* <0.01, \*\*\**P* <0.001. **(C)** Dominant TFs within each individual dataset defined by the top 10 upregulated (yellow) and bottom 10 downregulated (green) TFs based on activity scores.

Despite this diversity, transcriptional features consistently altered in at least two or three models revealed reproducible regulatory programs (**Fig. 3B**). Notably, resistant tumors displayed convergent epigenetic reprogramming characterized by the concurrent suppression of lineage-specifying factors *HNF4A* and *HNF4G*, alongside the downregulation of stress-responsive transcription factors *EGR1* and *KLF9*. Multiple models also showed activation of interferon signaling effectors *STAT2* and *IRF2*, reflecting parallel inflammatory signatures observed at the pathway level. The myeloid-lineage factor *SPI1* was also recurrently activated in H-NSCLC, M-CRC, and M-NSCLC, but not in the immune- and stroma-free M-BRCA model. In addition, proliferation-associated TFs including *E2F4* and its dimerization partner *TFDP1* were recurrently activated, aligning with the heightened cell-cycle activity characteristic of resistant tumors.

To dissect how these recurrent regulatory programs manifested within each respective tumor model, we examined the distinct regulatory profiles defined by the top and bottom 10 ranked TFs in each dataset (**Fig. 3C**). In H-NSCLC, the regulatory profile predominantly exhibited the hyper-proliferative phenotype characterized by the extensive activation of *MYC*, *E2F* and the metastasis-associated factor *FOXM1*. The suppression of *HNF4A* was particularly evident in M-NSCLC and M-CRC, where loss of epithelial identity is strongly manifested. Within the dataset-specific top-ranked TF profiles, *SPI1* was particularly prominent in H-NSCLC and M-NSCLC. In contrast, *STAT2* was recurrently observed in the resistance profiles of M-CRC and M-BRCA. However, the associated interferon-driven regulatory programs differed between the two models. Specifically, M-CRC exhibited coordinated involvement of *STAT2* with *IRF2* and *IRF1*, whereas M-BRCA preferentially engaged *IRF9*, consistent with a sustained interferon-responsive inflammatory state. Overall, these master regulators reflect the transcriptional circuitry that perpetuates the inflammatory–proliferative paradox and enforces the progressive loss of lineage identity.

### Gene-level executor modules consolidate the inflammatory–proliferative resistance program

Subsequently, we delineated gene expression signatures associated with these regulatory shifts by profiling transcriptomic changes recurrent across the four models. By focusing on differentially expressed genes (DEGs) that were consistently altered in at least three datasets, we identified conserved molecular effectors that functionally define the resistant phenotype (**Fig. 4A**). Among these shared genes, the upregulated signature was characterized by the co-occurrence of cell-cycle regulators, including *CDC7*, *NDC80* and *NCAPG*, together with immune/stress-associated genes such as *PTPN2*, *IFI16* and *TRIM59*. In contrast, a robust downregulated signature was marked by the suppression of lineage-specifying markers and structural components, exemplified by *NOTCH3*, *L1CAM*, and *CTSH*.

**Figure 4.**
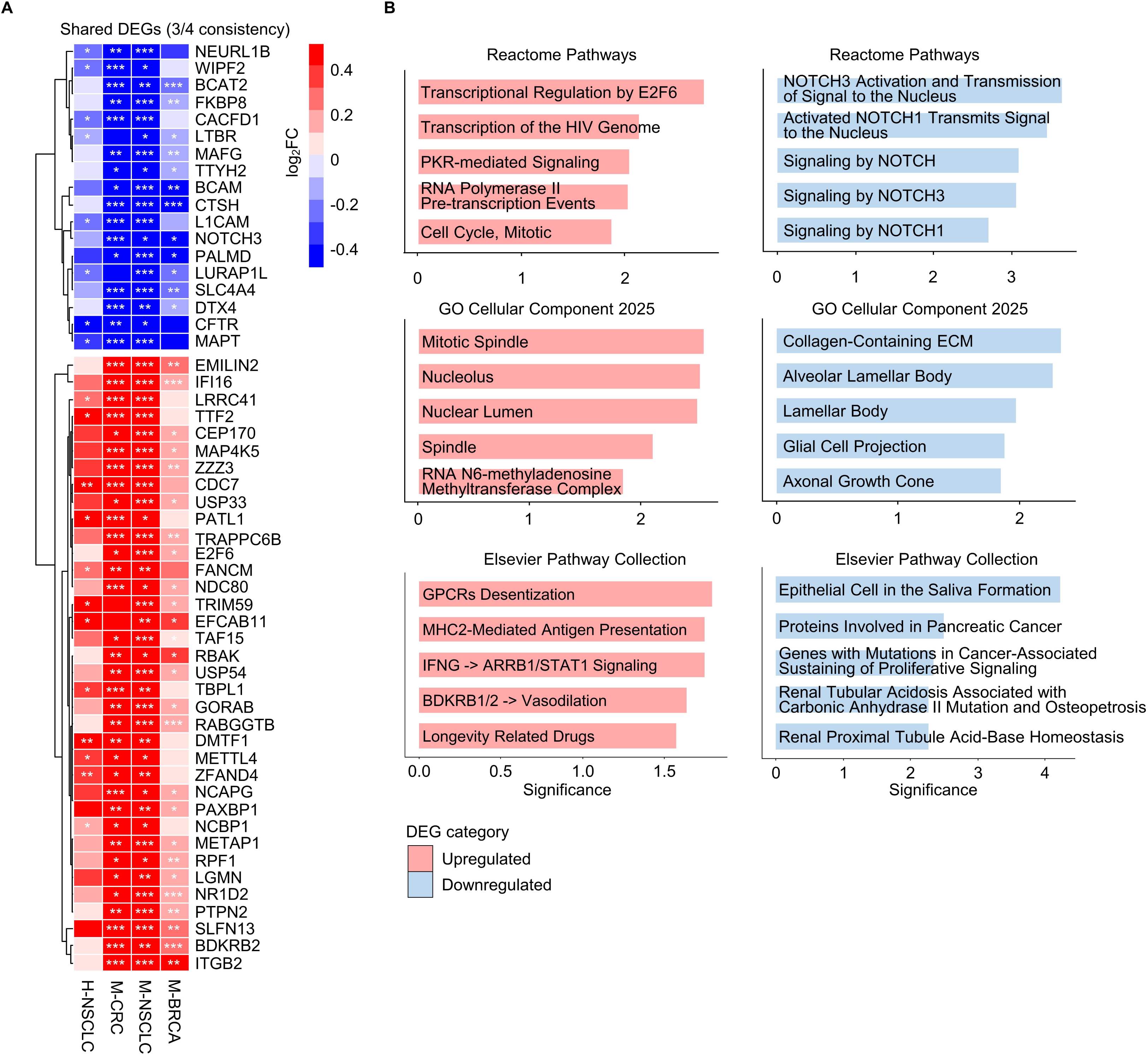
Conserved executor gene modules defining the acquired resistance phenotype. **(A)** Heatmap of differentially expressed genes (DEGs) consistently altered in at least three resistant models. The color scale represents the log_2_ fold-change (FC). \**P* <0.05, \*\**P* <0.01, \*\*\**P* <0.001. **(B)** Functional enrichment analysis of shared DEGs using “Reactome pathways 2024”, “GO cellular component 2025” and “Elsevier Pathway Collection” gene sets, displaying the top significantly enriched pathways for conserved upregulated (left) and downregulated (right) groups. Bar length indicates significance (-log_10_ *P*-value).

Pathway enrichment analysis further substantiated the biological implications of these shared signatures (**Fig. 4B**), based on over-representation analysis (ORA) across Reactome Pathways, GO Cellular Component, and Elsevier Pathway Collection. The upregulated DEGs were predominantly enriched in pathways related to cell-cycle regulation and interferon/innate immune–associated stress responses, including ‘Cell Cycle’ and ‘PKR-mediated signaling’. ORA also highlighted significant enrichment of ‘MHC2-Mediated Antigen Presentation’ and ‘IFNG to ARRB1/STAT1 Signaling’. In contrast, the downregulated DEGs reflected the loss of epithelial identity and tissue organization, evidenced by marked suppression of ‘Notch signaling’ and tissue-specific structural components such as ‘Alveolar Lamellar Body’ and ‘Collagen-containing Extracellular Matrix’. Collectively, these analyses indicate that acquired resistance across heterogeneous models is associated with a reproducible transcriptomic landscape featuring attenuation of lineage-associated structural pathways alongside concurrent activation of proliferative and immune/interferon-associated stress responses. Together, these gene-level patterns suggest separable functional themes that are subsequently resolved into higher-order modules by integrative network analysis.

### Integrated Pathway–TF–Gene network resolves a unified resistance program

To connect pathway-level convergence with its upstream regulators and downstream effectors, we integrated the recurrent signals from GSEA, TF activity inference, and shared DEGs into a unified resistance network (**Fig. 5A**). This network consistently organized into three tightly connected modules: hyper-proliferative outgrowth, active adaptation, and identity loss. The structural integrity of this modular organization was further validated by topological analysis, which revealed substantially higher internal than external connectivity across all three modules (internal density, 0.142–0.158; external density, 0.0059–0.0159; **Fig. S3**).

**Figure 5.**
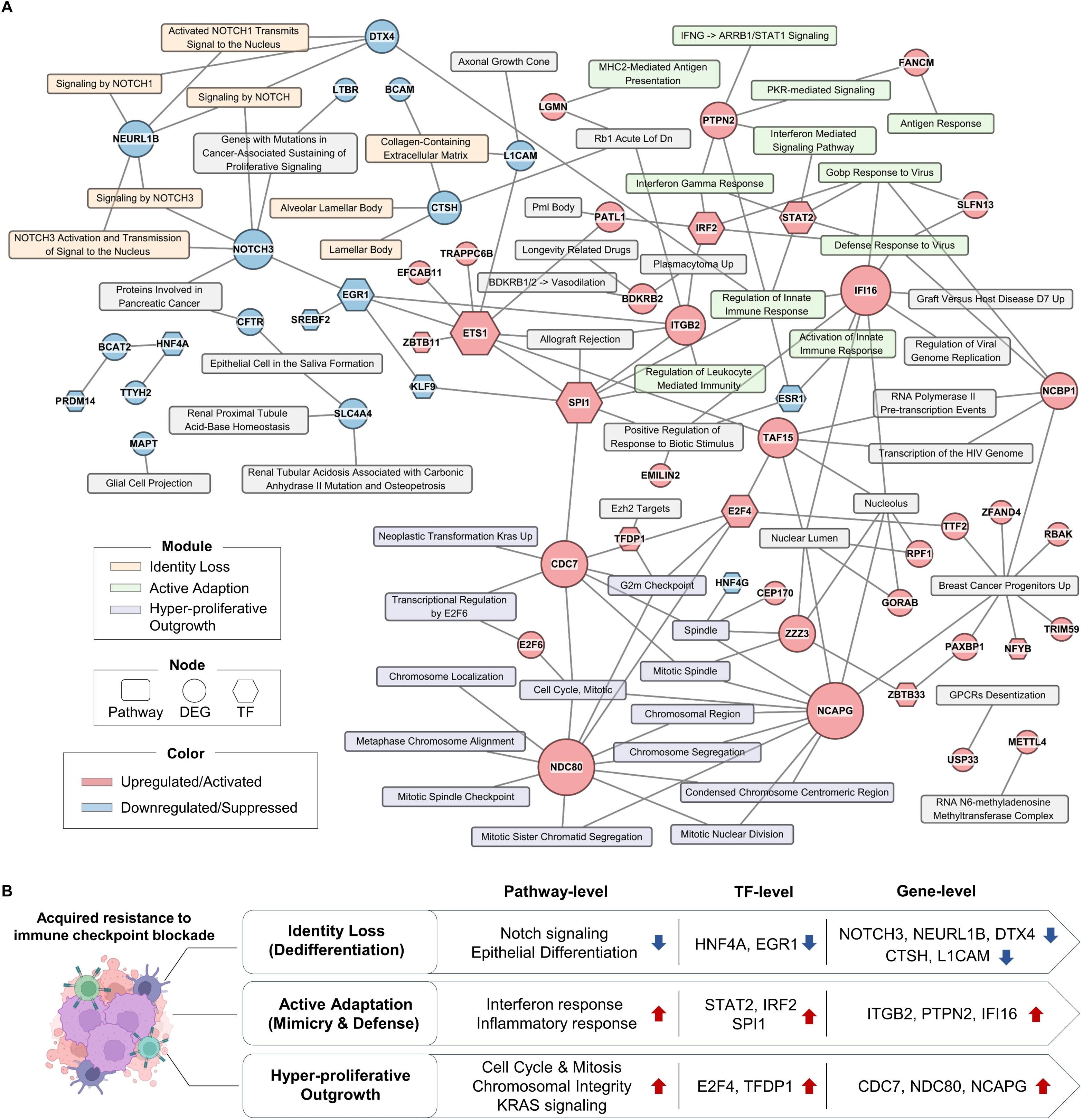
Integrated network analysis for a unified program of acquired resistance. **(A)** Multi-layer integration of recurrent signals across the four acquired resistance models. Nodes represent enriched pathways (rounded rectangles), transcription factors (TF; hexagons) and differentially expressed genes (DEG; circles). TF and DEG nodes are colored by activity score and log_2_FC, respectively, reflecting the direction of regulation, whereas pathway nodes are colored by functional module. Node size for TFs and DEGs is proportional to their connectivity (degree). Edges indicate curated links between layers. The integrated network resolves three major modules: Hyper-proliferative outgrowth, Active adaptation, and Identity loss. **(B)** Conceptual model summarizing the integrated network as a unified three-axis program of acquired resistance, in which identity loss, active adaptation (mimicry/defense), and hyper-proliferative outgrowth form a reinforcing system that stabilizes the resistant state.

The hyper-proliferative outgrowth module was centered on *E2F4* and *TFDP1*, coupled to replication and mitotic machinery genes such as *CDC7*, *NDC80*, and *NCAPG*, and linked to enrichment of cell-cycle and chromosome segregation programs including G2M checkpoint, mitotic spindle, and chromosome segregation gene sets. Notably, this proliferative axis also connected to KRAS-driven tumorigenic programs, consistent with aggressive neoplastic transformation during resistance. In parallel, the active adaptation module captured persistent inflammatory signaling together with defensive rewiring. This module was anchored by interferon-associated regulators *STAT2* and *IRF2*, and unexpectedly included activation of the myeloid regulator *SPI1*, coupled to effector genes such as *PTPN2*, *IFI16*, and *ITGB2*, and to enrichment of interferon gamma response, innate immune response, and response to virus pathways. Finally, the identity loss module featured lineage-specific remodeling involving the downregulation of NOTCH signaling–associated genes, *NOTCH3*, *NEURL1B*, and *DTX4*, with *EGR1* identified as a regulatory factor of this axis. These regulatory signatures were accompanied by structural components such as *CTSH* and *L1CAM*, reflecting a state of dedifferentiation and lineage plasticity under therapeutic pressure.

Together, these modules were not isolated features but were interconnected within a single network architecture, suggesting that acquired resistance involves coordinated coupling of inflammatory adaptation with proliferative acceleration and erosion of lineage constraints. Based on this integrated structure, we summarize acquired resistance as a unified three-axis tumor-cell-associated resistance program in which identity loss, active adaptation, and hyper-proliferative outgrowth form a reinforcing system that stabilizes the resistant state (**Fig. 5B**).

To assess the temporal dynamics of this three-axis program, we leveraged the multi-generational structure of the M-CRC dataset (CPar, C2nd, C4th). Across all three analytical layers, a coherent temporal hierarchy emerged. In the intermediate C2nd tumors, proliferative remodeling and lineage identity reprogramming were already apparent at the pathway, transcription factor, and gene expression levels. By contrast, immune and inflammatory adaptation became more distinctly activated in C4th (**Fig. S4**). These findings suggest that proliferative and identity-related changes arise earlier, while immune/inflammatory adaptation becomes more firmly established at later stages of resistance evolution.

## Discussion

Mechanistic inference of acquired ICB resistance has been constrained by reliance on single tumor models or platform-specific observations [17]. In this study, we integrated heterogeneous models into a unified resistance architecture using a design that combines independent within-dataset analyses with an intersection-based consensus strategy to prioritize reproducible signals and reduce model-specific artifacts. This approach aligns with network medicine concepts that emphasize convergent, module-level perturbations rather than isolated components [18]. Across four independent models, resistant states converged on a conserved “inflammatory–proliferative paradox” structured into three coupled axes: active inflammatory adaptation, hyper-proliferative outgrowth, and lineage identity loss. This architecture reflects a convergence-based logic where distinct evolutionary trajectories arrive at common functional states [19], connecting observations previously reported in individual settings [11–13] into a unified framework.

Our findings refine the interpretation of inflammation in the setting of acquired resistance. IFN-γ-related gene signatures correlate with response to PD-1 blockade in pretreatment samples, supporting the hypothesis that transcriptionally inflamed tumors respond better to ICB [20]. However, IFN signaling is context-dependent: prolonged IFN signaling can drive multigenic resistance programs through the upregulation of T-cell inhibitory receptor ligands [21], and chronic IFN exposure can establish epigenomic remodeling that sustains T-cell dysfunction even after the initial stimulus subsides [22]. Our results demonstrate that tumors with acquired resistance maintain robust IFN-γ response and innate immune signaling signatures across all four models, yet this inflammation no longer translates into effective tumor clearance. The concurrent reinforcement of cell-cycle and mitotic programs indicates that inflamed transcriptional states can be repurposed to support continued growth. In this reframing, inflammation in acquired resistance represents a stress-adaptive layer that coexists with, and may stabilize, proliferative outgrowth, rather than serving as a marker of productive antitumor immunity, consistent with the updated cancer-immunity cycle framework [23].

The inflammatory–proliferative program persisted not only in immune-competent tissue models (H-NSCLC and M-CRC) but also in settings with limited or absent non-tumor cellular components (M-NSCLC and M-BRCA). The persistence of interferon-associated signatures in the EMT6 model, which was established through chronic anti-PD-L1 exposure in the absence of immune cells, supports a tumor-cell-associated component of inflammatory adaptation. This interpretation is consistent with evidence for autocrine type I IFN signaling in cancer cells [24] and cell-autonomous regulation of resistance programs [25]. In the organoid-derived M-NSCLC model, where resistance was acquired under immune pressure *in vivo* but transcriptomes were profiled after *ex vivo* culture, the retention of inflammatory signatures is likewise consistent with a durable intrinsic component. More broadly, therapy pressure can select for, or induce, stable drug-tolerant states with transcriptional memory, as described in chromatin-mediated drug tolerance models [26]. Notably, the temporal pattern observed in M-CRC, in which proliferative remodeling and lineage identity reprogramming were already evident at the intermediate stage whereas immune/inflammatory adaptation became more pronounced at the terminal resistant stage, further supports a stepwise evolutionary process rather than an abrupt state transition. This pattern is compatible with models in which early proliferative and identity-related changes establish a permissive state upon which immune/inflammatory adaptation becomes progressively reinforced during resistance evolution.

We note, however, that compositional effects in the tissue-derived datasets cannot be fully excluded, and distinguishing clonal selection from stable reprogramming will require perturbation and longitudinal studies.

TF activity inference using regulon-based approaches identified a conserved regulatory backbone linking the inflammatory and proliferative programs. The IFN-associated regulators *STAT2* and *IRF2* were consistently activated. This pattern aligns with the reported duality of STAT2, in which early responses can support antitumor immunity, whereas sustained activation is associated with immunosuppression and chemoresistance [27]. IRF2 activity has also been implicated in IFN-mediated CD8+ T-cell exhaustion [28]. In parallel, *E2F4* and its dimerization partner *TFDP1* were convergently activated, providing a regulatory bridge from inflammatory signaling to cell-cycle progression; E2F4 has been increasingly recognized as an oncogenic driver promoting proliferation and therapy resistance [29]. The concurrent suppression of the lineage-specifying factors HNF4A and EGR1 completes the regulatory triad: HNF4A loss is associated with dedifferentiation and lineage plasticity in epithelial cancers [30], and EGR1 downregulation is associated with therapy resistance [31]. Together, these patterns are compatible with the broader view that lineage plasticity represents a shared route to resistance across cancer types [32]. Finally, the activation of *SPI1* (PU.1), a canonical myeloid-lineage factor [33], in the three models with prior in vivo immune exposure (H-NSCLC, M-CRC, and M-NSCLC), but not in the immune- and stroma-free M-BRCA system, points to a predominantly microenvironment-associated origin. In the ex vivo M-NSCLC model, where resistance was acquired under immune pressure in vivo and transcriptomes were profiled after organoid culture, a residual tumor-associated contribution cannot be excluded; however, distinguishing tumor-intrinsic immune mimicry from persistent myeloid influence will require single-cell or spatial validation.

At the effector gene level, conserved differentially expressed genes provide testable nodes operationalizing the three-axis architecture. In the hyper-proliferative axis, recurrent upregulation of *CDC7*, *NDC80*, and *NCAPG* is consistent with aggressive cell-cycle reinforcement. *CDC7*, a replication kinase essential for origin firing, has emerged as a therapeutic target whose inhibition exposes replication stress vulnerabilities [34]. *NDC80* and *NCAPG*, involved in kinetochore function and chromosome condensation respectively, have been associated with tumorigenesis and poor prognosis [35, 36]. In the inflammatory adaptation axis, *IFI16* participates in STING-dependent innate sensing [37], while *PTPN2*, identified by *in vivo* CRISPR screening as an immunotherapy target [38], is now targeted by ABBV-CLS-484, a first-in-class inhibitor in clinical evaluation [39]. ITGB2 has emerged as a functionally relevant mediator of inflammatory adaptation that promotes immunosuppressive microenvironmental remodeling by attenuating anti-tumor immune activation [40, 41]. In the identity loss axis, the coordinated downregulation of NOTCH3 and its trafficking regulators, NEURL1B and DTX4, may indicate attenuation of Notch signaling, potentially weakening lineage constraint and favoring dedifferentiation-associated tumor progression in a context-dependent manner [42]. CTSH is expressed in branching bronchial epithelium during lung development and is required for normal pulmonary surfactant processing, underscoring its association with epithelial lineage programs [43]. Similarly, suppression of L1CAM has been associated with a more stem-like state in pancreatic cancer, suggesting its potential role in driving dedifferentiation [44]. We note that some effector genes are immune-lineage associated and may reflect compositional effects. These associations are hypothesis-generating and will require functional validation.

Translationally, our results caution against interpreting inflammatory signatures as unequivocal indicators of ICB responsiveness in the acquired resistance setting [11]. Because the three modules are interconnected, single-axis targeting may be insufficient, motivating combination strategies [45] that address inflammatory adaptation together with proliferative and plasticity-associated programs. Consistent with this logic, preclinical studies support combining ICB with CDK4/6 inhibitors that enhance antigen presentation and activate endogenous retroviral programs [46], as well as epigenetic modulators that can reverse chronic IFN-associated chromatin states [22]. Because inflammatory adaptation is coupled to proliferative and lineage-associated programs, our framework provides a rationale for multi-node interventions, including polypharmacological natural products, as candidate approaches to modulate multiple resistance-associated pathways [47, 48]. Transcriptome-informed frameworks can further support the systematic evaluation of such multi-target agents [49–51] and their immunomodulatory potential as ICB combination partners [52].

An important consideration is the heterogeneity of resistance definitions across models. Our cross-model integration is intended to identify conserved resistance-associated programs rather than to equate the biological meaning or effect sizes of resistance across systems. Although anti–PD-1 and anti–PD-L1 agents are not mechanistically identical, the convergence across both therapies suggests shared downstream selection pressures under chronic inhibition of the PD-1/PD-L1 axis, consistent with broadly comparable clinical activity reported across multiple settings [53]. In the immune-free M-BRCA system, part of the adaptation may reflect tumor-cell–associated responses to PD-L1 ligation, consistent with reports of cell-autonomous PD-L1 signaling [54, 55]. Quantitative differences across models should therefore be interpreted cautiously as context-dependent, motivating follow-up validation in harmonized experimental systems and longitudinal clinical cohorts.

Several limitations warrant consideration. First, our framework is based on reanalysis of public bulk transcriptomic datasets without experimental perturbation, and therefore the proposed architecture is hypothesis-generating rather than causal. Second, bulk profiling cannot fully disentangle tumor-state regulation from microenvironmental composition, particularly for immune-lineage-associated regulons, and cell-type-resolved approaches will be important for validation [56]. Third, the murine models relied on surrogate anti-mouse PD-1 antibodies, which differ in binding properties from clinically approved agents such as nivolumab and pembrolizumab that do not cross-react with murine PD-1 [57], potentially influencing quantitative estimates of shared signals. To address these gaps, we have established a PBMC-humanized mouse model that will enable evaluation under clinically approved human anti-PD-1 treatment in future studies [58]. Future investigations using this platform, combined with single-cell and spatial transcriptomic profiling [59, 60], will be critical for validating the identified resistance modules in a human-relevant immune context.

In conclusion, cross-species transcriptomic integration of four acquired ICB resistance models reveals a conserved inflammatory-proliferative paradox organized into three interconnected modules. This architecture challenges the binary hot/cold paradigm by demonstrating that inflammatory signaling in resistant tumors reflects tumor-cell-associated adaptation rather than productive antitumor immunity. The regulatory backbone linking *STAT2*/*IRF2*, *E2F4*/*TFDP1*, and *HNF4A*/*EGR1* provides tractable nodes for therapeutic intervention. Our framework prioritizes combination strategies that simultaneously address inflammatory adaptation, proliferative outgrowth, and lineage plasticity to restore durable immunotherapy responses.

## Acknowledgments

This work was supported by grants from the National Research Foundation of Korea (RS-2023-00210923), Basic Science Research Program through the National Research Foundation of Korea funded by the Ministry of Education (RS-2025-25397599).

## Declaration of interest statement

The authors declare no conflicts of interest.

## Notes

https://www.ncbi.nlm.nih.gov/geo/query/acc.cgi?acc=GSE248249

https://www.ncbi.nlm.nih.gov/geo/query/acc.cgi?acc=GSE249000

https://www.ncbi.nlm.nih.gov/geo/query/acc.cgi?acc=GSE261889

https://www.ncbi.nlm.nih.gov/geo/query/acc.cgi?acc=GSE186034

### Competing Interest Statement

The authors have declared no competing interest.

## References

1. Sharma, P. and J.P. Allison, The future of immune checkpoint therapy. Science, 2015. 348(6230): p. 56–61.

2. Dong, W., et al., The mechanism of anti–pd-l1 antibody efficacy against pd-l1–negative tumors identifies nk cells expressing pd-l1 as a cytolytic effector. Cancer discovery, 2019. 9(10): p. 1422–1437.

3. Topalian, S.L., et al., Mechanism-driven biomarkers to guide immune checkpoint blockade in cancer therapy. Nature Reviews Cancer, 2016. 16(5): p. 275–287.

4. Zhang, N., et al., Biomarkers and prognostic factors of PD-1/PD-L1 inhibitor-based therapy in patients with advanced hepatocellular carcinoma. Biomarker research, 2024. 12(1): p. 26.

5. Sharma, P., et al., Primary, adaptive, and acquired resistance to cancer immunotherapy. Cell, 2017. 168(4): p. 707–723.

6. Jenkins, R.W., D.A. Barbie, and K.T. Flaherty, Mechanisms of resistance to immune checkpoint inhibitors. British journal of cancer, 2018. 118(1): p. 9–16.

7. Hanahan, D. and R. Weinberg, Hallmarks of cancer: the next generation. cell. 2011 Mar 4; 144 (5): 646–74.

8. Karasaki, T., et al., Evolutionary characterization of lung adenocarcinoma morphology in TRACERx. Nature medicine, 2023. 29(4): p. 833–845.

9. Chen, K., T.W. Shuen, and P.K. Chow, The association between tumour heterogeneity and immune evasion mechanisms in hepatocellular carcinoma and its clinical implications. British Journal of Cancer, 2024. 131(3): p. 420–429.

10. Wu, B., et al., Cold and hot tumors: from molecular mechanisms to targeted therapy. Signal transduction and targeted therapy, 2024. 9(1): p. 274.

11. Memon, D., et al., Clinical and molecular features of acquired resistance to immunotherapy in non-small cell lung cancer. Cancer Cell, 2024. 42(2): p. 209–224.e9.

12. Shi, Y., et al., Acquired resistance to PD-L1 inhibition enhances a type I IFN-regulated secretory program in tumors. EMBO reports, 2024. 26(2): p. 521.

13. Wang, M., et al., Acquired resistance to immunotherapy by physical barriers with cancer cell–expressing collagens in non–small cell lung cancer. Proceedings of the National Academy of Sciences, 2025. 122(24): p. e2500019122.

14. Kuleshov, M.V., et al., Enrichr: a comprehensive gene set enrichment analysis web server 2016 update. Nucleic acids research, 2016. 44(W1): p. W90–W97.

15. Badia-i-Mompel, P., et al., decoupleR: ensemble of computational methods to infer biological activities from omics data. Bioinformatics advances, 2022. 2(1): p. vbac016.

16. Garcia-Alonso, L., et al., Benchmark and integration of resources for the estimation of human transcription factor activities. Genome research, 2019. 29(8): p. 1363–1375.

17. Restifo, N.P., M.J. Smyth, and A. Snyder, Acquired resistance to immunotherapy and future challenges. Nature Reviews Cancer, 2016. 16(2): p. 121–126.

18. Barabási, A.-L., N. Gulbahce, and J. Loscalzo, Network medicine: a network-based approach to human disease. Nature reviews genetics, 2011. 12(1): p. 56–68.

19. Konieczkowski, D.J., C.M. Johannessen, and L.A. Garraway, A convergence-based framework for cancer drug resistance. Cancer cell, 2018. 33(5): p. 801–815.

20. Ayers, M., et al., IFN-γ–related mRNA profile predicts clinical response to PD-1 blockade. The Journal of clinical investigation, 2017. 127(8): p. 2930–2940.

21. Benci, J.L., et al., Opposing functions of interferon coordinate adaptive and innate immune responses to cancer immune checkpoint blockade. Cell, 2019. 178(4): p. 933–948.e14.

22. Qiu, J., et al., Cancer cells resistant to immune checkpoint blockade acquire interferon-associated epigenetic memory to sustain T cell dysfunction. Nature cancer, 2023. 4(1): p. 43–61.

23. Mellman, I., et al., The cancer-immunity cycle: Indication, genotype, and immunotype. Immunity, 2023. 56(10): p. 2188–2205.

24. Cheon, H., et al., How cancer cells make and respond to interferon-I. Trends in cancer, 2023. 9(1): p. 83–92.

25. Benci, J.L., et al., Tumor interferon signaling regulates a multigenic resistance program to immune checkpoint blockade. Cell, 2016. 167(6): p. 1540–1554.e12.

26. Sharma, S.V., et al., A chromatin-mediated reversible drug-tolerant state in cancer cell subpopulations. Cell, 2010. 141(1): p. 69–80.

27. Canar, J., et al., The duality of STAT2 mediated type I interferon signaling in the tumor microenvironment and chemoresistance. Cytokine, 2023. 161: p. 156081.

28. Lukhele, S., et al., The transcription factor IRF2 drives interferon-mediated CD8+ T cell exhaustion to restrict anti-tumor immunity. Immunity, 2022. 55(12): p. 2369–2385.e10.

29. Matsubara, J., et al., The E2F4 transcriptional repressor is a key mechanistic regulator of colon cancer resistance to irinotecan (CPT-11). bioRxiv, 2025.

30. Cammareri, P., et al., Loss of colonic fidelity enables multilineage plasticity and metastasis. Nature, 2025. 644(8076): p. 547–556.

31. Baron, V., et al., The transcription factor Egr1 is a direct regulator of multiple tumor suppressors including TGFβ1, PTEN, p53, and fibronectin. Cancer gene therapy, 2006. 13(2): p. 115–124.

32. Mehta, A. and B.Z. Stanger, Lineage plasticity: the new cancer hallmark on the block. Cancer research, 2024. 84(2): p. 184–191.

33. Tenen, D.G., et al., Transcription factors, normal myeloid development, and leukemia. Blood, The Journal of the American Society of Hematology, 1997. 90(2): p. 489–519.

34. Quintanal-Villalonga, A., et al., CDC7 inhibition impairs neuroendocrine transformation in lung and prostate tumors through MYC degradation. Signal Transduction and Targeted Therapy, 2024. 9(1): p. 189.

35. Tooley, J. and P.T. Stukenberg, The Ndc80 complex: integrating the kinetochore’s many movements. Chromosome research, 2011. 19(3): p. 377–391.

36. Sun, H., et al., NCAPG promotes the oncogenesis and progression of non-small cell lung cancer cells through upregulating LGALS1 expression. Molecular cancer, 2022. 21(1): p. 55.

37. Jønsson, K., et al., IFI16 is required for DNA sensing in human macrophages by promoting production and function of cGAMP. Nature communications, 2017. 8(1): p. 14391.

38. Manguso, R.T., et al., In vivo CRISPR screening identifies Ptpn2 as a cancer immunotherapy target. Nature, 2017. 547(7664): p. 413–418.

39. Baumgartner, C.K., et al., The PTPN2/PTPN1 inhibitor ABBV-CLS-484 unleashes potent anti-tumour immunity. Nature, 2023. 622(7984): p. 850–862.

40. Paierhati, P., B. Ma, and M. Abudukeremu, Exosomal ITGB2 Mediates Immune Evasion in Triple-Negative Breast Cancer by Suppressing Dendritic Cell Activation via TLR4. Iranian Journal of Public Health, 2025. 54(6): p. 1252.

41. Li, C., et al., Identifying ITGB2 as a potential prognostic biomarker in ovarian cancer. Diagnostics, 2023. 13(6): p. 1169.

42. Ku, S.-Y., et al., Notch signaling suppresses neuroendocrine differentiation and alters the immune microenvironment in advanced prostate cancer. The Journal of Clinical Investigation, 2024. 134(17).

43. Bühling, F., et al., Expression of cathepsins B, H, K, L, and S during human fetal lung development. Developmental dynamics: an official publication of the American Association of Anatomists, 2002. 225(1): p. 14–21.

44. Cave, D.D., et al., TGF-β1 secreted by pancreatic stellate cells promotes stemness and tumourigenicity in pancreatic cancer cells through L1CAM downregulation. Oncogene, 2020. 39(21): p. 4271–4285.

45. Patel, S.A. and A.J. Minn, Combination cancer therapy with immune checkpoint blockade: mechanisms and strategies. Immunity, 2018. 48(3): p. 417–433.

46. Goel, S., et al., CDK4/6 inhibition triggers anti-tumour immunity. Nature, 2017. 548(7668): p. 471–475.

47. Yeo, H., S.-Y. Kim, and S.-M. Park, Harnessing transcriptomics for discovery of natural products to overcome acquired cancer resistance. Archives of Pharmacal Research, 2025: p. 1–29.

48. Lee, C., Phytochemicals as promising agents in Axl-targeted cancer treatment. The Korean Journal of Physiology & Pharmacology: Official Journal of the Korean Physiological Society and the Korean Society of Pharmacology, 2025. 29(5): p. 533–545.

49. Park, M., et al., KORE-Map 1.0: Korean medicine Omics Resource Extension Map on transcriptome data of tonifying herbal medicine. Scientific data, 2024. 11(1): p. 974.

50. Seo, E.-H., et al., KORE-Map 1.1: Korean medicine omics resource extension map on transcriptome data of dyspepsia herbal medicine. BMC Genomic Data, 2026.

51. Baek, S.-J., et al., Multidimensional transcriptome dataset for systematic evaluation of Jakyakgamcho-tang-induced cell signatures. Scientific Data, 2026.

52. Kim, S.-Y., et al., Deciphering the immunomodulatory mechanisms of Bojungikki-tang via systematic transcriptomic and immune cell interaction network analysis. Biomedicine & Pharmacotherapy, 2025. 188: p. 118129.

53. Zhou, L., et al., Efficacy and safety of first-line PD-1/PD-L1 inhibitors combined with or without anti-angiogenesis therapy for extensive-stage small-cell lung cancer: a network meta-analysis. Therapeutic advances in medical oncology, 2025. 17: p. 17588359251348310.

54. Clark, C.A., et al., Tumor-intrinsic PD-L1 signals regulate cell growth, pathogenesis, and autophagy in ovarian cancer and melanoma. Cancer research, 2016. 76(23): p. 6964–6974.

55. Gato-Cañas, M., et al., PDL1 signals through conserved sequence motifs to overcome interferon-mediated cytotoxicity. Cell reports, 2017. 20(8): p. 1818–1829.

56. Lauss, M., et al., Molecular patterns of resistance to immune checkpoint blockade in melanoma. Nature communications, 2024. 15(1): p. 3075.

57. Schofield, D.J., et al. Activity of murine surrogate antibodies for durvalumab and tremelimumab lacking effector function and the ability to deplete regulatory T cells in mouse models of cancer. in MAbs. 2021. Taylor & Francis.

58. Chuprin, J., et al., Humanized mouse models for immuno-oncology research. Nature Reviews Clinical Oncology, 2023. 20(3): p. 192–206.

59. Aung, T.N., et al., Spatial signatures for predicting immunotherapy outcomes using multi-omics in non-small cell lung cancer. Nature Genetics, 2025. 57(10): p. 2482–2493.

60. Lee, M.Y., et al., Single-cell spatial transcriptomics uncovers niches that govern response to PD-1/PD-L1 blockade in cutaneous squamous cell carcinoma. Journal for Immunotherapy of Cancer, 2026. 14(1): p. e014067.

