## Supplementary figure for "Transcriptomic Integration Reveals a Conserved Inflammatory–Proliferative Paradox in Acquired Resistance to Immune Checkpoint Blockade"

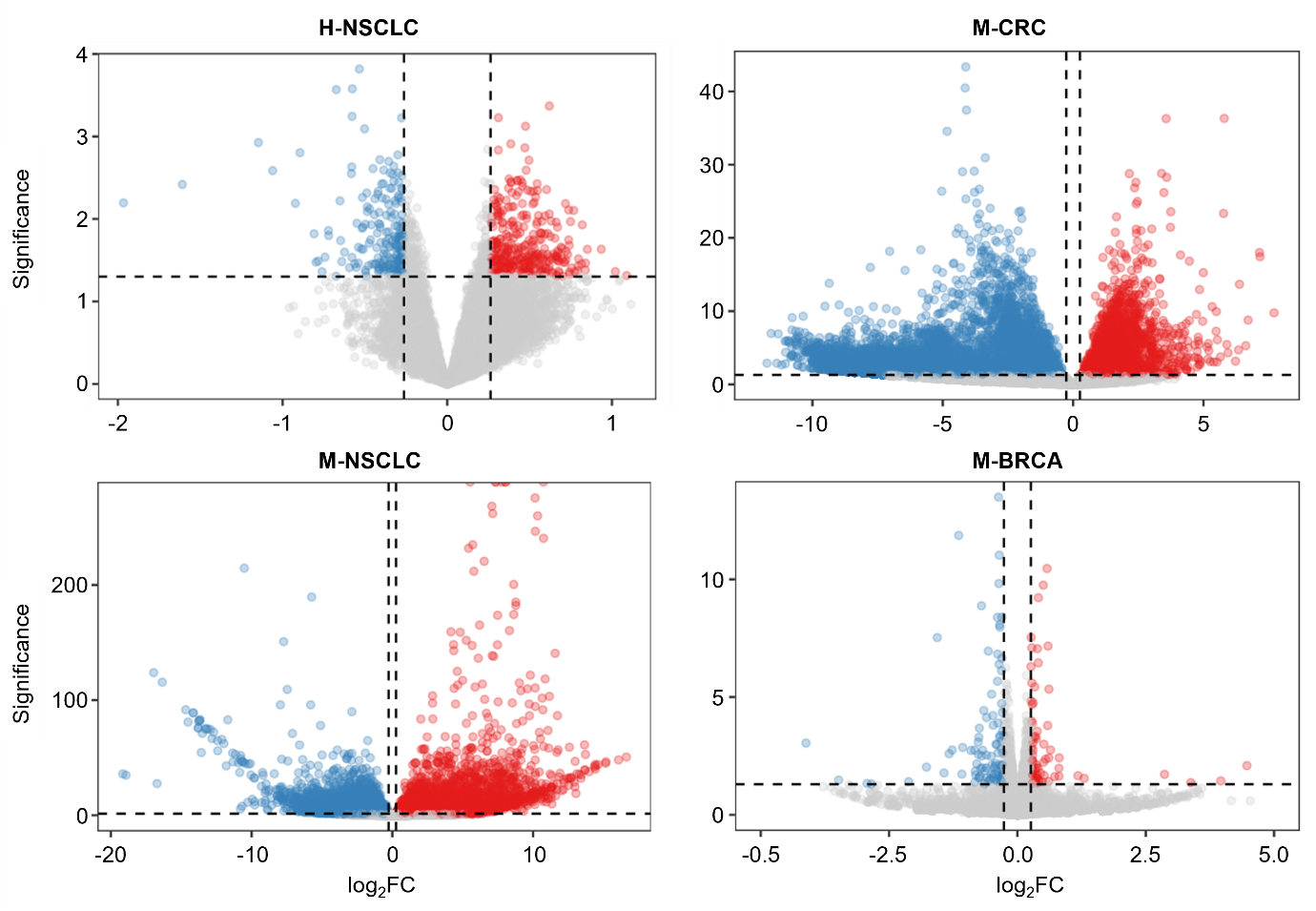

**Figure S1. Volcano plots for four acquired resistance models.** Differentially expressed genes in H-NSCLC, M-CRC, M-NSCLC, M-BRCA are shown as volcano plots. The x-axis represents log_2_ fold-change (FC), and the y-axis indicates significance (-log_10_ *P*-value). Red and blue dots represent significantly upregulated and downregulated genes, respectively, defined by |FC| ≥ 1.2 and *P* < 0.05. Gray dots indicate non-significant genes. Dashed lines mark the FC and *P*-value thresholds.

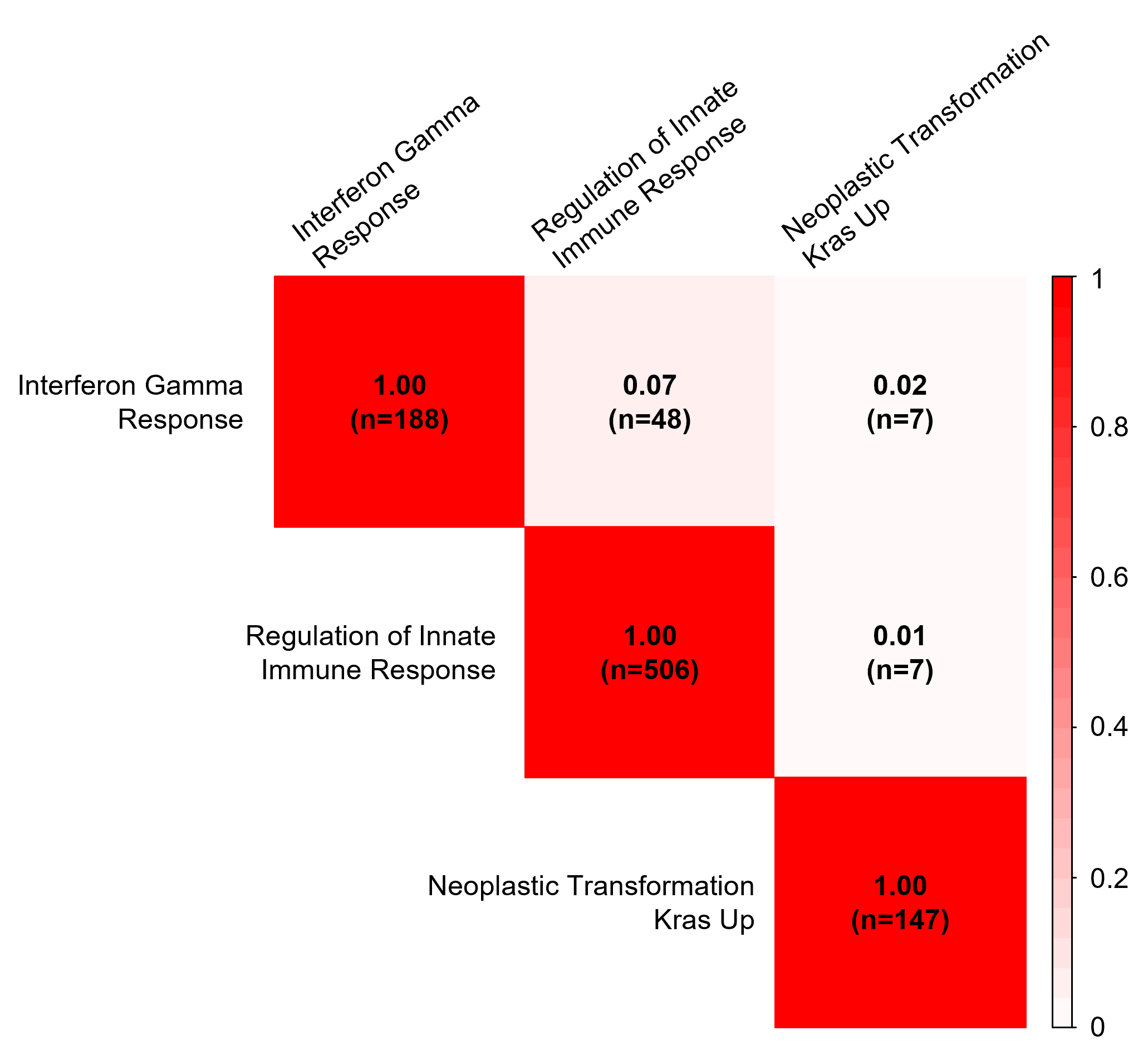

**Figure S2. Pairwise Jaccard similarity among three pathways consistently regulated across four datasets.**

The pairwise overlap between the gene sets of three pathways was quantified using the Jaccard index and visualized as a heatmap. Values within each cell represent the Jaccard index. The value 'n' indicates the total number of genes in each pathway (diagonal) or the number of shared genes between two pathways (off-diagonal). Low off-diagonal Jaccard values (0.01–0.07) indicate high specificity of each gene set with minimal inter-pathway overlap.

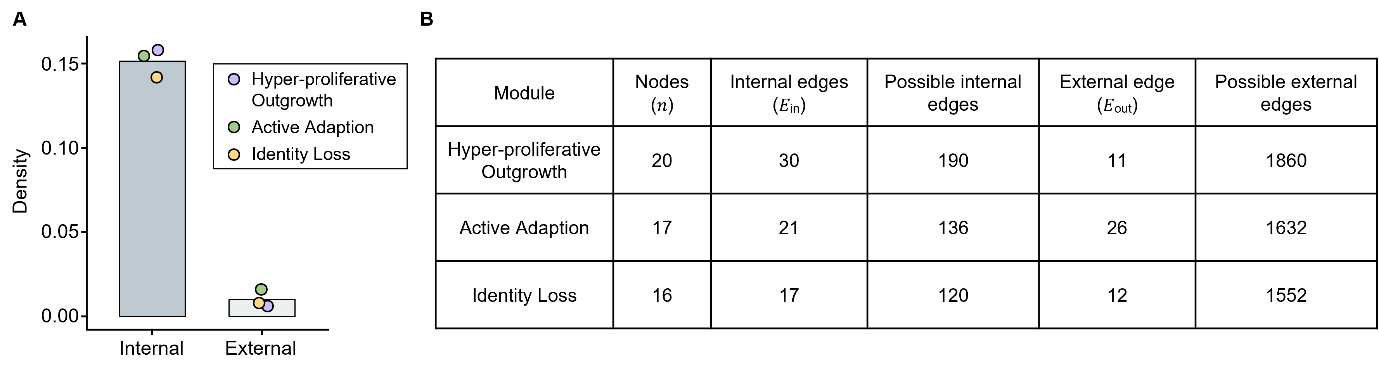

**Figure S3. Topological characteristics of the three functional modules within the integrated network.**

The integrated network (𝑁 = 113 nodes, 𝐸 = 158 edges) consists of three functional modules with distinct connectivity profiles. **(A)** Comparison of internal and external densities across the three modules. Bars represent the mean density, and colored points indicate the values for individual modules. Internal refers to interactions within each module, while external denotes interactions between module components and nodes in the surrounding network. All three modules showed substantially higher internal than external density, indicating that each forms a relatively cohesive subnetwork. **(B)** Summary of network topology metrics for each module, including the number of nodes, observed internal and external edges, and the theoretical maximum of internal and external edges used for density calculation.

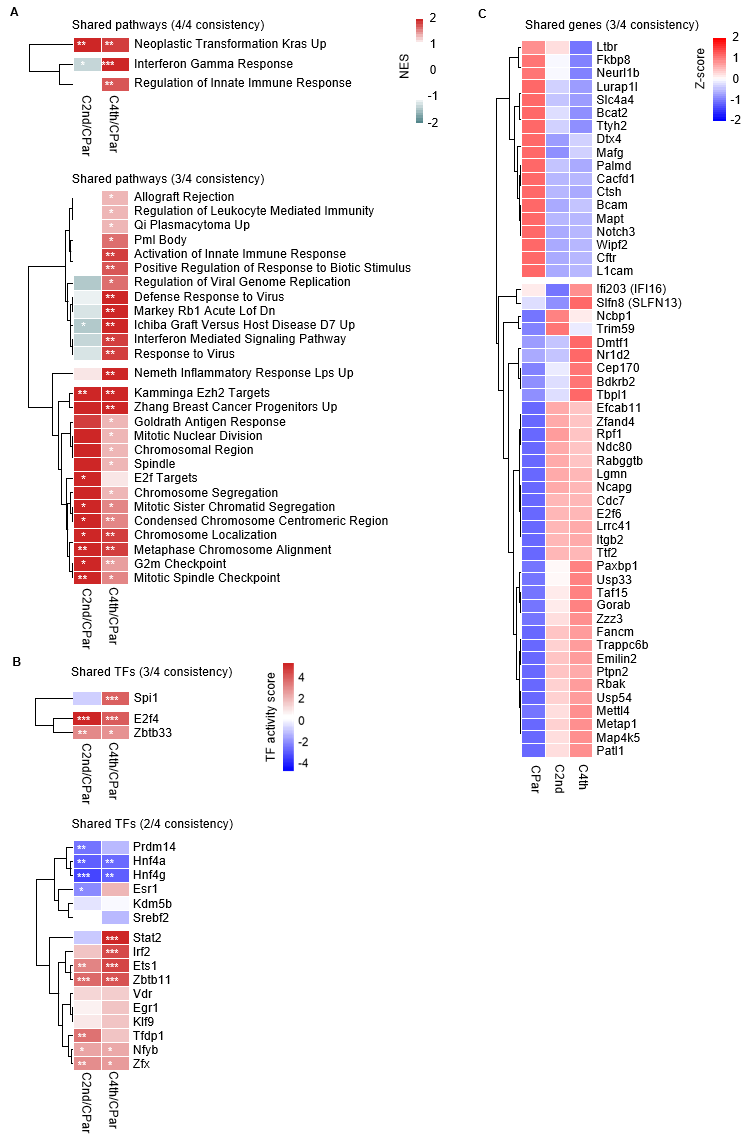

**Figure S4. Stage-wise dynamics of the three-axis resistance program in the M-CRC model.** Consistent molecular features identified from four resistant datasets were assessed across the parental (CPar), second-generation resistant (C2nd), and fourth-generation resistant (C4th) states. **(A)** Pathway-level changes in C2nd and C4th relative to CPar, represented by of normalized enrichment scores (NES), are shown for pathways selected based on cross-dataset consistency. *FDR < 0.05, **FDR < 0.01, ***FDR < 0.001. **(B)** Transcription factor (TF)-level alterations in C2nd and C4th relative to CPar, represented by TF activity scores, are shown for TFs selected based on cross-dataset consistency. **P* < 0.05, ***P* < 0.01, ****P* < 0.001. **(C)** Gene-level expression patterns in CPar, C2nd, and C4th, shown as z-score values for genes consistently altered in at least three datasets.
